# Understanding the effects of oxytocin receptor variants on OXT–OXT receptor binding: A mathematical model

**DOI:** 10.1101/2024.02.28.582600

**Authors:** Preeti Dubey, Yingye Fang, K. Lionel Tukei, Shobhan Kuila, Xinming Liu, Annika Sahota, Antonina I. Frolova, Erin L. Reinl, Manasi Malik, Sarah K. England, Princess I. Imoukhuede

## Abstract

Approximately half of U.S. women giving birth annually receive Pitocin, the synthetic form of oxytocin (OXT), yet its effective dose can vary significantly. This variability presents safety concerns due to unpredictable responses, which may lead to adverse outcomes for both mother and baby. To address the need for improved dosing, we developed a data-driven mathematical model to predict OXT receptor (OXTR) binding. Our study focuses on five prevalent OXTR variants (V45L, P108A, L206V, V281M, and E339K) and their impact on OXT–OXTR binding dynamics in two distinct cell types: human embryonic kidney cells (HEK293T), commonly used in experimental systems, and human myometrial smooth muscle cells, containing endogenous OXTR. We parameterized the model with cell-specific OXTR surface localization measurements. To strengthen the robustness of our study, we conducted a comprehensive meta-analysis of OXT– OXTR binding, enabling parameterization of our model with cell-specific OXT–OXTR binding kinetics (myometrial OXT–OXTR K_d_ = 1.6 nM, kon = 6.8 × 10^5^ M^−1^ min^−1^, and koff = 0.0011 min^−1^). Our meta-analysis revealed significant homogeneity in OXT–OXTR affinity across experiments and species with a K_d_ = 0.52 - 9.32 nM and mean K_d_ = 1.48 ± 0.36 nM. Our model achieves several valuable insights into designing dosage strategies. First, we predicted that the OXTR complex reaches maximum occupancy at 10 nM OXT in myometrial cells and at 1 µM in HEK293T cells. This information is pivotal for guiding experimental design and data interpretation when working with these distinct cell types, emphasizing the need to consider effects for specific cell types when choosing OXTR-transfected cell lines. Second, our model recapitulated the significant effects of genetic variants for both experimental and physiologically relevant systems, with V281M and E339K substantially compromising OXT–OXTR binding capacity. These findings suggest the need for personalized oxytocin dosing based on individual genetic profiles to enhance therapeutic efficacy and reduce risks, especially in the context of labor and delivery. Third, we demonstrated the potential for rescuing the attenuated cell response observed in V281M and E339K variants by increasing the OXT dosage at specific, early time points. Cellular responses to OXT, including Ca^2+^ release, manifest within minutes. Our model indicates that providing V281M- and E339K-expressing cells with doubled OXT dose during the initial minute of binding can elevate OXT–OXTR complex formation to levels comparable to wild-type OXTR. In summary, our study provides a computational framework for precision oxytocin dosing strategies, paving the way for personalized medicine.

## INTRODUCTION

Oxytocin (OXT) is a peptide hormone that regulates a multitude of physiological functions within the brain and reproductive systems. Specifically, OXT plays a crucial role in parturition and lactation, as well as in regulating several social and cognitive functions, including mating, aggression, and fear^1^. Recent evidence also suggests the involvement of the OXT system in neuropsychiatric conditions, such as autism spectrum and anxiety-related disorders^2^. Importantly, the synthetic form of OXT, Pitocin, is given to nearly two million women annually to induce or augment labor in the USA^3^. However, the recommended range of OXT dose varies widely, by up to 20-fold^4–6^. Higher doses of OXT lead to a high risk of uterine rupture and hyperstimulation, whereas lower doses may prolong labor, leading to fetal infection and the need for cesarean delivery^5–9^. The serious consequences of inappropriate OXT dosing have motivated research on understanding and predicting the differences in OXT responsiveness amongst individuals.

Further complicating the determination of appropriate OXT dosing, OXT–OXT receptor (OXTR) interactions vary between cell types, and the OXT dose response in biologically relevant cells remains inadequately explored. For instance, specific naturally occurring genetic variants, such as V281M and E339K, in the *OXTR* gene significantly hamper the surface localization of OXTRs, resulting in reduced OXT–OXTR binding and diminished OXT response in HEK293T cells overexpressing exogenous OXTR^5^. However, HEK293T cells lack the endogenous machinery for OXTR trafficking and signaling. Yet, challenges in transfecting the biologically relevant cells, human myometrial smooth muscle cells, have prevented these experimental studies. Additionally, in a clinical setting, access to myometrial cells in order to test OXT sensitivity is not feasible. A mathematical model that accurately predicts appropriate OXT dosing in biologically relevant cells and tissues would have important implications for maternal health during labor and delivery. The parameterization of such a model should take into consideration both the known influence of dysregulated OXTR surface localization on OXT–OXTR interaction and the biological relevance of the experimental models from which all data are derived.

The current study addresses this gap by constructing an OXT–OXTR binding framework that incorporates the cell type-specific association rate (k_on_) and dissociation rate (k_off_) of OXT–OXTR binding and OXTR variant-specific concentrations on the surface of HEK293T cells or human myometrial smooth muscle cells. Our model offers predictions on dose-dependent OXT-OXTR binding dynamics, particularly in the myometrial cells where traditional experimental approaches are limited. Our model also provides the first insights into how the heterologous system, HEK293T cells, differs from a biologically relevant context (myometrial cells). Such differential insights are essential for enhancing the biological relevance and limitations of the data obtained from heterologous systems.

## RESULTS

### A mathematical model for predicting the effects of genetic variants on OXT–OXTR binding

Five of the most prevalent naturally occurring genetic variants, P108A, V45L, E339K, L206V, and V281M (**Figure 1**), alter OXTR activation and signaling in HEK293T cells.^5^ We constructed a mathematical model to better understand the effects of these genetic variants on OXT–OXTR binding dynamics. Specifically, this model predicts the OXT–OXTR binding and complex formation elicited by the OXT concentrations used in *in vitro* experiments.^5^ The key feature of the model is its OXTR-specific and cell-specific parameterization. First, the model uses distinctly different cell-surface OXTR concentrations precisely measured for each of the five variants (**Table 1**). Second, the model uses cell-type specific association rates (k_on_) and dissociation rates (k_off_) of OXT-OXTR interaction, because the values differ by cell type, species, and even gestational time (**Table 2**). These parametrizations are described in detail below.

**Figure 1:**
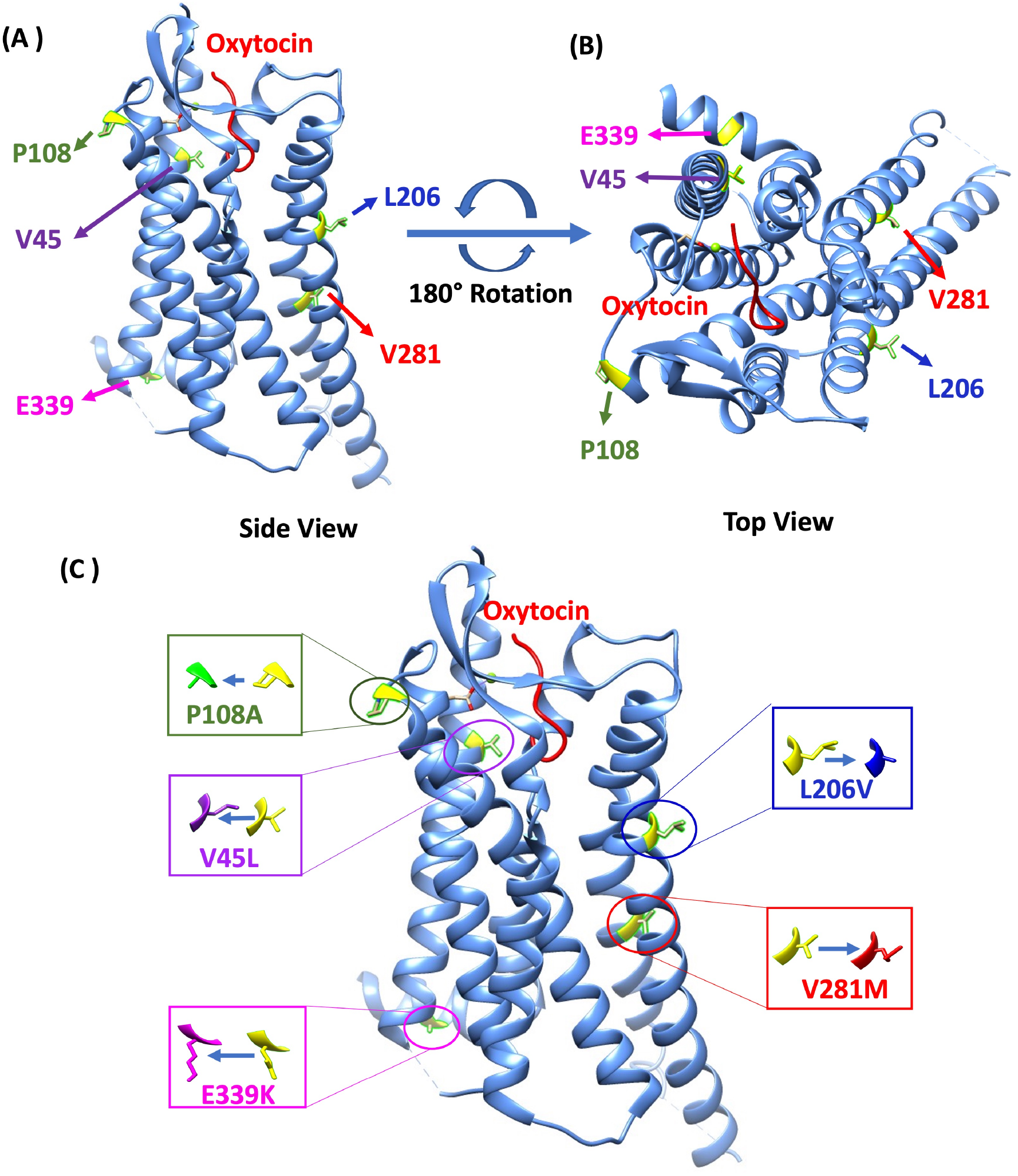
Schematic of oxytocin-bound OXTR structure with mutation sites. **(A)** A visual representation of the OXTR and oxytocin ligand as well as the specific locations of mutation sites. The image depicts a side-view perspective, parallel to the plane of the plasma membrane. The OXTR protein and the hormone oxytocin are shown as ribbon-like structures. **(B)** A top view of the OXTR/oxytocin complex as observed from the extracellular space. **(C)** Variant amino acid residue substitutions and their locations in the crystal structure of OXTR (PDB: 7RYC), generated using UCSF Chimera software. The various color-coded boxes correspond to several OXTR variants: blue denotes the L206V variant, red denotes the V281M variant, magenta denotes the E339K variant, purple denotes the V45L variant, and green denotes the P108A variant.

**Table 1:**
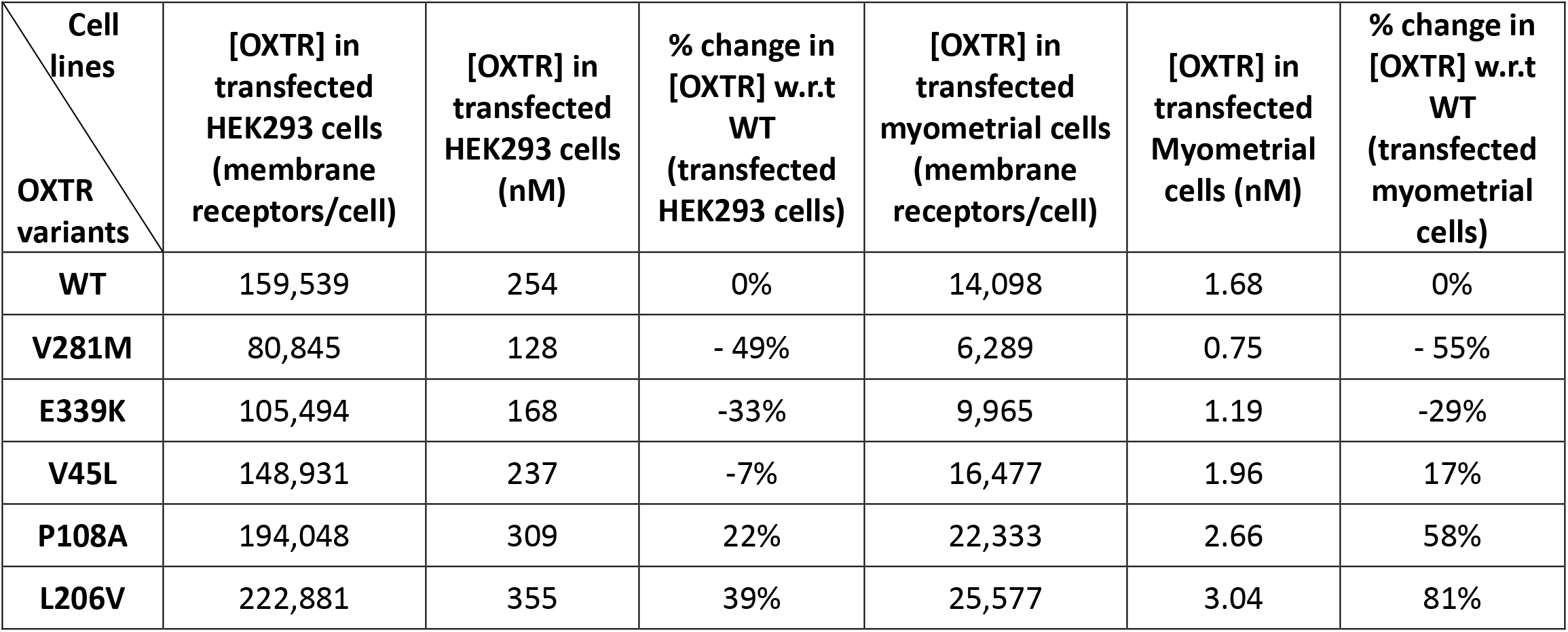
Wild type and variant OXTR receptor concentration on cell surface in copies/cell and in nM.

| Cell lines<br>OXTR variants | [OXTR] in transfected HEK293 cells (membrane receptors/cell) | [OXTR] in transfected HEK293 cells (nM) | % change in [OXTR] w.r.t WT (transfected HEK293 cells) | [OXTR] in transfected myometrial cells (membrane receptors/cell) | [OXTR] in transfected Myometrial cells (nM) | % change in [OXTR] w.r.t WT (transfected myometrial cells) |
| --- | --- | --- | --- | --- | --- | --- |
| WT | 159,539 | 254 | 0% | 14,098 | 1.68 | 0% |
| V281M | 80,845 | 128 | - 49% | 6,289 | 0.75 | - 55% |
| E339K | 105,494 | 168 | -33% | 9,965 | 1.19 | -29% |
| V45L | 148,931 | 237 | -7% | 16,477 | 1.96 | 17% |
| P108A | 194,048 | 309 | 22% | 22,333 | 2.66 | 58% |
| L206V | 222,881 | 355 | 39% | 25,577 | 3.04 | 81% |

**Table 2:**
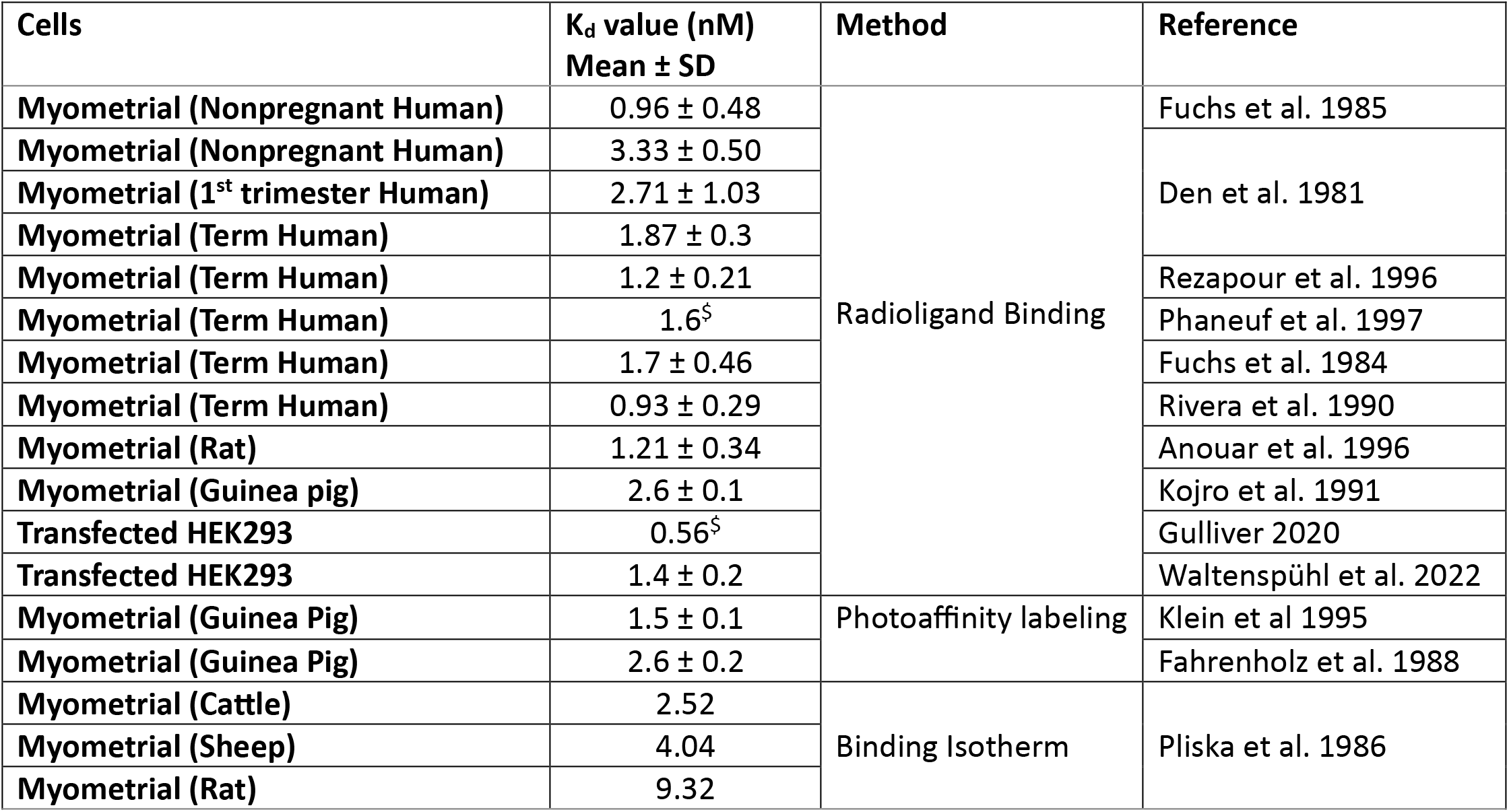
Reported binding affinities of OXTR in different species and cell lines. SD is the standard deviation. ^$^ The values used in the model.

### Cell-specific and OXTR–specific parameterization: differential OXTR surface localization by genetic variants

In HEK293T cells, the differential surface localization of OXTR genetic variants is associated with altered responses to OXT^5^. However, to understand the impact of these genetic variants on OXTR surface localization in biologically relevant myometrial cells, further investigation is needed to inform the development of the mathematical model. In myometrial cells, we found that the V281M and E339K variants significantly reduced the proportion of OXTR on the cell surface, whereas the P108A and L206V variants increased OXTR on the cell surface (**Table 1**). These results were similar to our findings, and those reported by others^12^, in HEK293T cells. We found a differential response in HEK293T and myometrial cells for V45L that displayed subtle differences in surface OXTR expression with 7% fewer OXTR molecules on HEK293T cells and 17% more OXTR on myometrial cells, when compared to wilt type (WT) OXTR (**Table 1**). The number of OXTR molecules on HEK293T cells^5^ and human myometrial cells^10^ were measured using quantitative flow cytometry and converted to molar concentrations (**Table 1**).

### Cell-specific and OXTR-specific parameterization: OXT–OXTR binding affinities

To accurately represent the OXT–OXTR binding dynamics in our model, we performed an extensive literature survey of OXT-OXTR binding affinity (K_d_, K_d_ = k_off_/ k_on_) and conducted a meta-analysis on the literature mined K_d_ values. We found that K_d_ values of OXT–OXTR binding range from 0.56 nM up to 9.32 nM, varying with the measurement methods, species, and cell types (**Table 2 and Figure 2**). For instance, we observed that OXTRs in human myometrium at term (≥37 weeks gestation) bind to OXT with greater affinity (lower K_d_), compared to ‘non-pregnant’ and ‘1^st^ trimester’ myometrial OXTRs. Importantly, the choice of measurement method can lead to several-fold variations in K_d_ values. For example, rat myometrial K_d_ is 1.21 nM by radioligand binding and 9.32 nM by binding isotherm analysis (**Table 2**).

**Figure 2:**
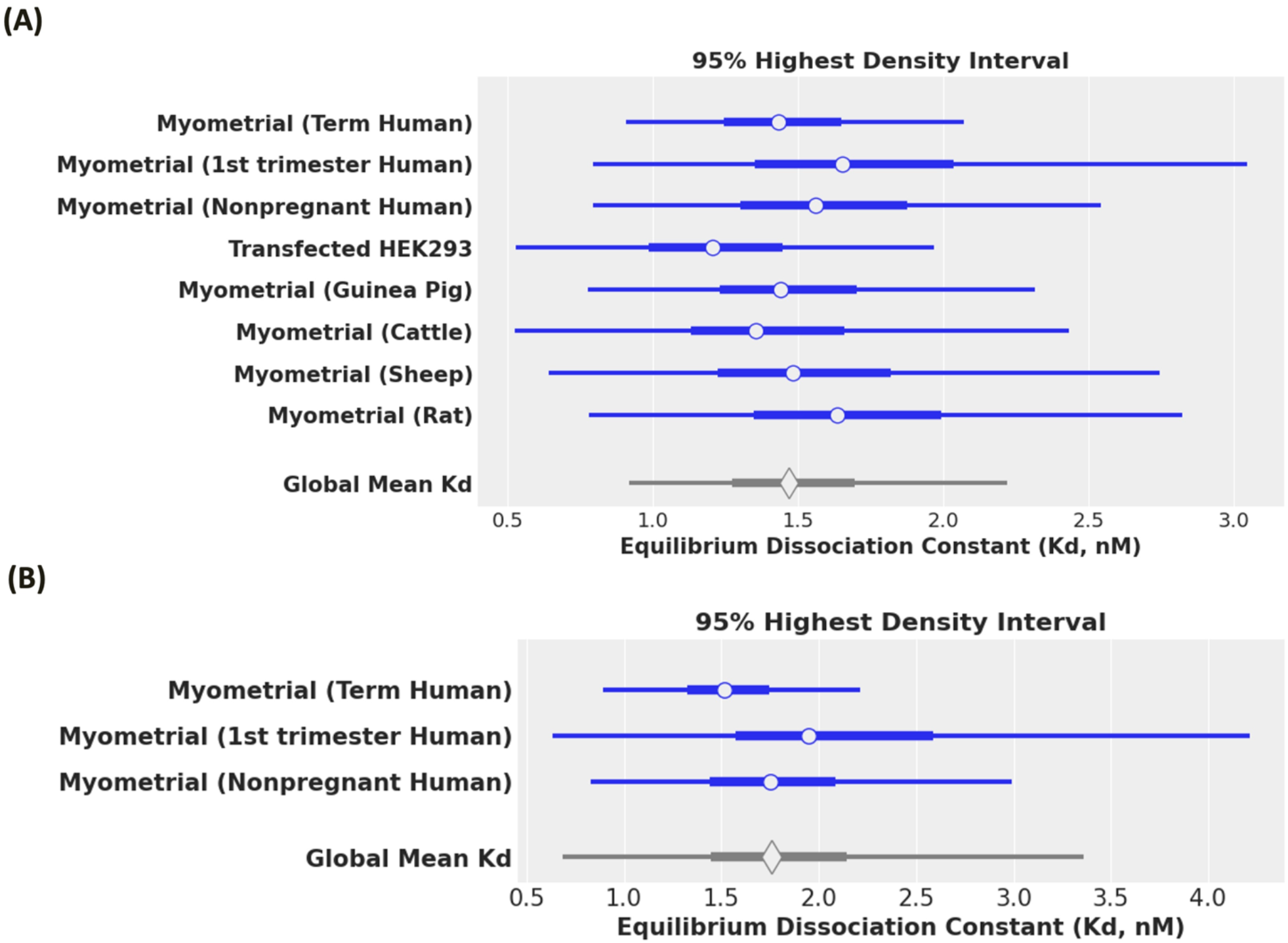
OXT–OXTR binding affinities differ based on species, cell type, and gestational status. High density interval (HDI) forest plot of equilibrium dissociation constant (K_d_) for oxytocin-to-oxytocin receptor (OXTR) binding in different cell lines and species. **(A)** The cells included in the figure are myometrial cells from term human, 1^st^ trimester human, nonpregnant human, guinea pig, cattle, sheep, rat and HEK293 cells transfected to express the OXTR. The blue horizontal lines represent the 95% HDI, with the thicker segment indicating the credible interval (the 95% probable values of the parameter). The blank circle marker shows the mean. The black vertical line shows the global mean of equilibrium dissociation constant, K_d_ = 1.50 nM with 95% HDI [0.92, 2.22] 95% credible interval [1.27, 1.70]. **(B)** The myometrial cells from humans only (term human, 1^st^ trimester human, and nonpregnant human) were shown in the HDI forest plot of equilibrium dissociation constant (K_d_) for oxytocin-to-oxytocin receptor (OXTR) binding. The blue horizontal lines represent the 95% HDI, with the thicker segment indicating the credible interval (the 95% probable values of the parameter). The blank circle marker shows the mean. The black vertical line shows the global mean of equilibrium dissociation constant, K_d_ = 1.78 nM with 95% HDI [0.68, 3.36] and 95% credible interval [1.45, 2.14].

With this variation in K_d_, we needed to determine the value to use for parametrization in the myometrial and HEK293 models. While the variation in K_d_ values for human myometrial cells varied from 0.93 nM to 3.33 nM, only the Phaneuf *et al*. ^11^ study provided sufficient data for us to extract k_on_ and k_off_ values. Thus, we parameterized the myometrial model with a K_d_ of 1.6 nM. Notably, 1.6 nM is close to the representative K_d_ value across all studies (1.48 ± 0.51 nM) determined by our meta-analysis (**Figure 2, myometrial term human**). For HEK293 cells, we found two sources of K_d_ values, 0.56 nM^12^ and 1.4 nM^13^. In this study, we selected the K_d_ value of 0.56 nM for HEK293 cells because this value was measured using the same radioligand binding method as used in Phaneuf *et al*. for the myometrial cells^11^. Consequently, the OXT-OXTR binding affinity in human myometrial cells is threefold weaker than in HEK293T cells in our models. This difference in K_d_ values could be due to the distinct cellular characteristics of each cell type. Our human myometrial cell model and HEK293T cell model used different K_d_, k_on_, and k_off_ values: k_on_ = 6.8 × 10^5^ M^−1^ min^−1^ and k_off_ = 0.0011 min^−1^ (K_d_ = 1.6 nM) for human myometrial cells^11^ and k_on_ = 8.8 × 10^6^ M^−1^ min^−1^ and k_off_ = 0.005 min^−1^ (K_d_ = 0.56 nM) for HEK293T cells^12^ (**Table 2**). Our model assumes the binding kinetics, k_on_ and k_off_, are not affected by the genetic variants because none of these variants occur at the OXT binding site (**Figure 1**).

### Internal model validation: Half-maximal occupancy predictions consistent with K_d_

Because the binding affinity (K_d_=k_off_/k_on_) defines the concentration where equilibrium binding capacity is half-maximal, an internal model validation was conducted by simulating OXT-OXTR binding at OXT concentrations equivalent to the K**_d_** values for HEK293T cells ([OXT] = 0.56 nM) and human myometrial cells ([OXT] = 1.6 nM, **Figure 3 A & B**). The equilibrium was reached at 10 hours in HEK293T cells and at 36 hours in human myometrial cells. As expected, the number of bound receptors at equilibrium in HEK293T cells is ∼78,000 complexes/cell (∼50% of the free receptors, **Table 1**), and in myometrial cells, it is ∼7000 complexes/cell (∼50% of the free receptors, **Table 1**). Thus, our model is mathematically accurate.

**Figure 3:**
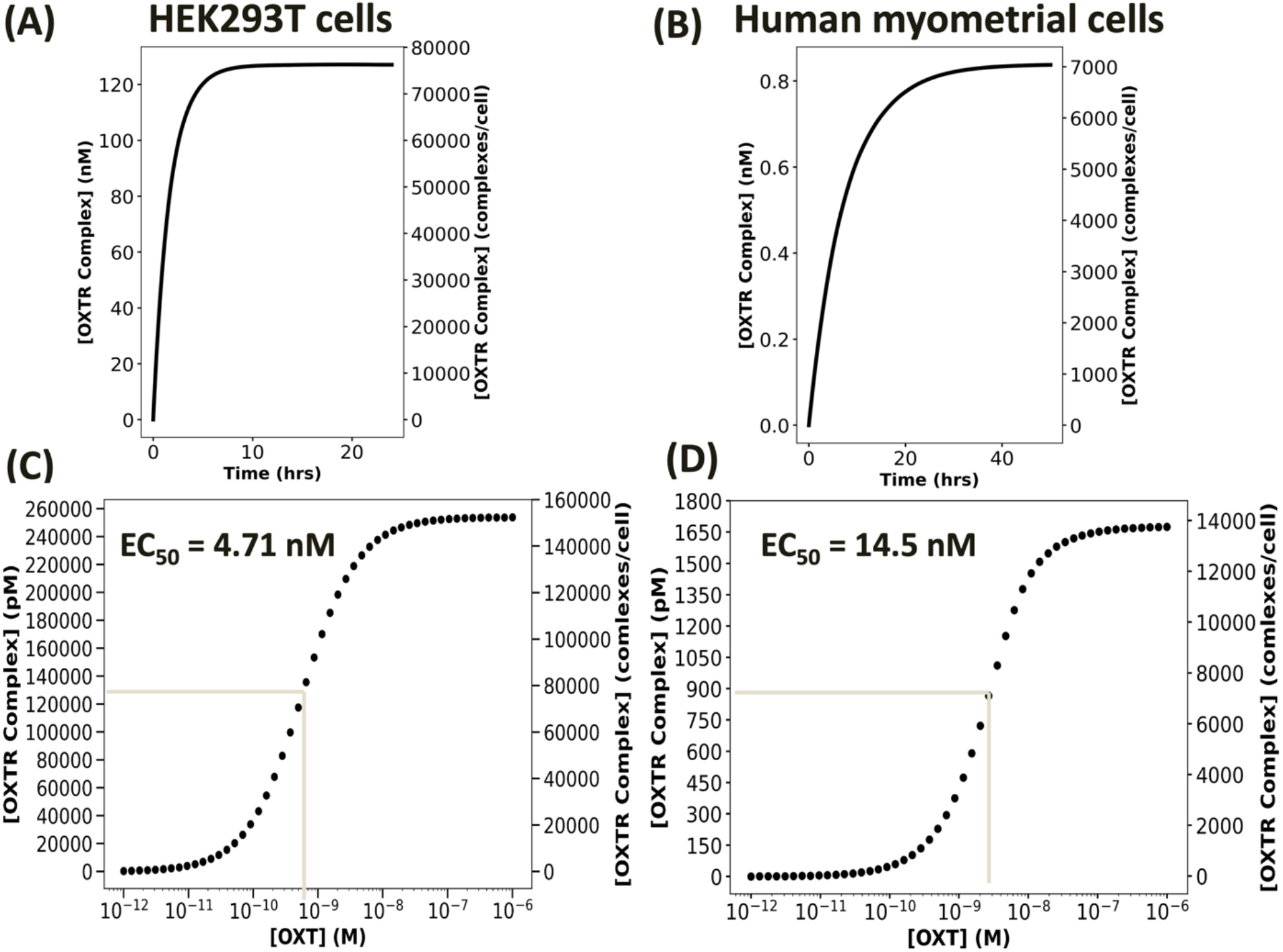
The mathematical model half-maximal occupancy predictions align with experimental K_d_ data for OXT-OXTR binding. Measures of wild-type OXTRC formation when [OXT] = Kd in **(A)** HEK293T cells (K_d_ = 0.56 nM) and **(B)** human myometrial cells (K_d_ = 1.6 nM). The initial concentration of wild-type OXTR in HEK293T cells was 254 nM (159,539 copies/cell), and in myometrial cells it was 1.68 nM (14,098 copies/cell). **Oxytocin dose response curve for (C)** HEK293 cells and **(D)** human myometrial cells. The left y-axis shows bound WT OXTR complex in pM and right Y-axis shows the bound WT OXTR complex in complexes/cell. The maximal WT OXTR complex formation was recorded at 10 hours for 50 increasing oxytocin doses ranging between 1pM to 1μM.

### Model validation to experimental data: OXT EC_50_ and OXTR occupancy predictions align with experimental observations

To illustrate that our model can recapitulate experimental outcomes, we first simulated the OXT dose-response curve at OXT concentrations varying from 1 pM to 1 µM and calculated an EC_50_ of 4.71 nM OXT for wild-type OXTR in HEK293T cells (**Figure 3C**). This EC_50_ of OXT-OXTR binding aligns with the EC_50_ of OXT-induced OXTR activation in HEK293T cells, which is 5.4 nM.^5^ Using this framework, we calculated that the EC_50_ of OXT-OXTR binding in human myometrial cells is 14.5 nM (**Figure 3D**), which also aligns with the experimentally measured EC_50_ for OXTR activation in human myometrial cells (30 nM).^14^

We further validated our model with three initial OXT concentrations (10 pM, 10 nM, and 1 µM) that have been used to induce OXTR activation in HEK293T cells.^5^ Our simulation of HEK293T cell binding dynamics demonstrates a strong alignment with the *in vitro* experimental observations of OXTR activation. In experimental data, 10 pM OXT is too low to form OXT–OXTR complexes (OXTRCs) in HEK293T cells (**Figure 4A**) and barely induces calcium flux responses.^5^ In HEK293T cells, 10 nM OXT forms 6000 WT OXTRCs/cell, which is only a fraction (∼4% of total surface OXTRs) of the maximal potential (**Figure 4B**). Nonetheless, 10 nM OXT induces ∼40% of maximal calcium flux in vitro,^5^ suggesting that 6000 WT OXTRCs/cell may be sufficient to induce measurable calcium flux. The highest dose, 1 µM OXT, induces the maximal OXTR activation, which is also seen experimentally as the maximal calcium flux.^5^

**Figure 4:**
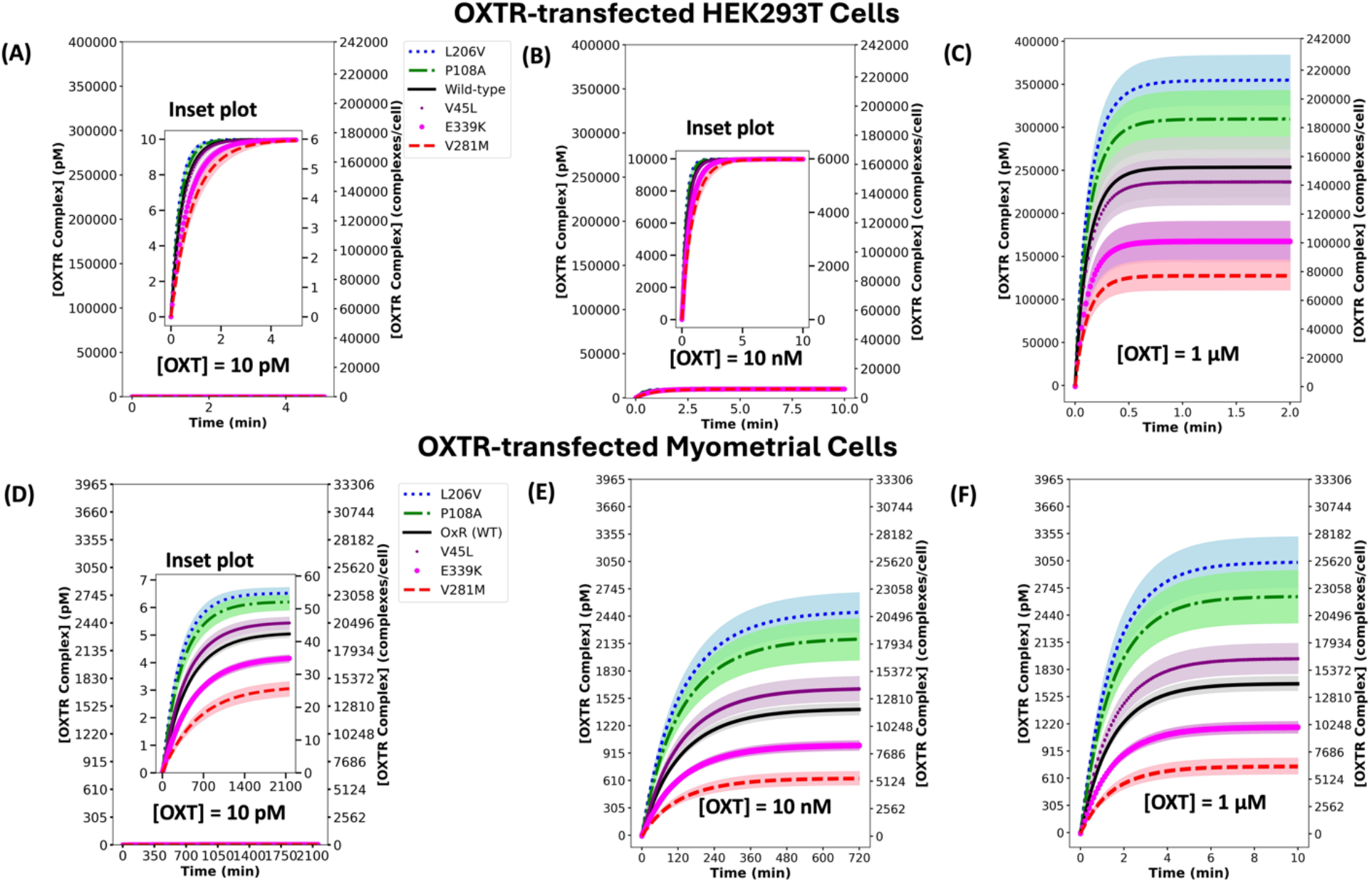
Simulation of HEK293T cell binding dynamics demonstrates a strong alignment with the *in vitro* experimental observations of OXTR activation. The OXTR complex (OXTRC) formation for all variants at equilibrium in OXTR-transfected **(A-C)** HEK293T cells and **(D-F)** human myometrial cells treated with 10 pM **(A & D)**, 10 nM **(B & E)**, and 1 μM **(C & F)** OXT treatments. The plot shows the concentration of bound wild-type (WT) OXTR and 5 variant OXTRs in pM on the left Y-axis and in complexes/cell on the right Y-axis. The zoomed in plot is shown as inset plot in figures **(A), (B) and (D).** The black solid line represents wild type, the blue dotted line represents variant L206V, the green dashed-dotted line represents variant P108A, the purple point–marked line represents variant V45L, the magenta circle–marked line represents variant E339K, and the red dashed line represents variant V281M. The initial concentrations used for OXTR on the surface of a myometrial cell were as follows: [WT] = 1.68 nM, [V281M] = 0.75 nM, [P108A] = 2.66 nM, [L206V] = 3.04 nM, [V45L] = 1.96 nM, [E339K] = 1.19 nM. The initial concentrations used for OXTR (nM) in HEK293T cells are as follows: [WT] = 254 nM, [V281M] = 128 nM, [P108A] = 309 nM, [L206V] = 355 nM, [V45L] = 237 nM, [E339K] = 168 nM.

### Model application: Human myometrial cells reach near-maximal OXTR occupancy at lower OXT doses than HEK293T cells

To compare OXT–OXTR binding in physiologically relevant cells with that of a heterologous system, we used our model to determine the binding dynamics of OXT–OXTR in myometrial cells at the same three initial OXT concentrations used to induce OXTR activation in HEK293T cells. As expected, human myometrial cells show near-maximum occupancy at 10 nM OXT, because this value is ∼10 fold greater than the reported human myometrial K_d_ (1.6 nM). Indeed, 10 nM OXT is almost as effective as 1 µM OXT for inducing the maximal OXTRC formation, ∼12,000 WT OXTRCs/cell (**Figure 4**E & F). However, at 10nM OXT requires ∼10 hours to reach near-maximal occupancy, while 1 µM OXT only takes 10 minutes. At 10 pM OXT, which is 150 times lower than the myometrial K_d_, the concentration is too low to form OXTRCs on myometrial cells. Fewer than 50 occupied WT OXTR complexes were predicted (**Figure 4D)**.

Compared with HEK293T cells, myometrial cells possess three times weaker OXT–OXTR binding affinity and express 10 times fewer WT OXTRs on the cell surface. However, myometrial cell OXTR can be adequately activated at a lower OXT concentration (10 nM vs 1 µM), but take 10 times longer to reach maximal OXTR occupancy at 1 µM OXT (10 min vs. 1 min, **Figure 4C vs.** 4F), compared to HEK293T cells.

### Model application: Predicting the effects of genetic variants on complex formation in myometrial cells at experimental OXT concentrations

The effect of genetic variants on OXT-OXTR binding in human myometrial cells has not been measured experimentally. Our model predicted the effects of genetic variants on OXT-OXTR complex formation in human myometrial cells at three OXT concentrations that were previously experimentally tested in HEK293T cells. At all concentrations, variants P108A and L206V showed an increase in OXTRC formation while variants V281M and E339K show a decrease in OXTRC formation as compared to wild type OXTR (**Figure 4D, 4E, and 4F**). While this effect was found at both 10 pM (**Figure 4D**) and 10 nM OXT (**Figure 4E**), the largest difference was seen at 1 µM OXT, which elicited the maximal OXTRC formation. Variants P108A and L206V showed an increase of 58% and 81% in OXTRC formation, while variants V281M and E339K showed a decrease of 55% and 29% in OXTRC formation.

OXTRC formation at the upper and lower bounds of OXTR initial concentration overlapped significantly for the two variants P108A and L206V in all three simulations (shaded blue and green regions in **Figures 4D-F**). This overlap suggests that these two variants exhibit comparable complex formation capacity.

### Strategizing OXT dosage and treatment duration to activate genetic variants with reduced OXTR activation in myometrial cells

We applied the model to test whether a high-dose regimen (e.g., > 1 µM OXT) could rescue the OXTRC formation for V281M and E339K OXTRs in myometrial cells at equilibrium. Our model shows even with high OXT doses, the OXTRC levels in V281M and E339K cells are ∼57% and ∼28% lower compared with the wild type at equilibrium (**Figures 5A, 6A**). This effect is likely due to the lower OXTR availability in V281M- and E339K-expressing myometrial cells relative to OXT.

**Figure 5:**
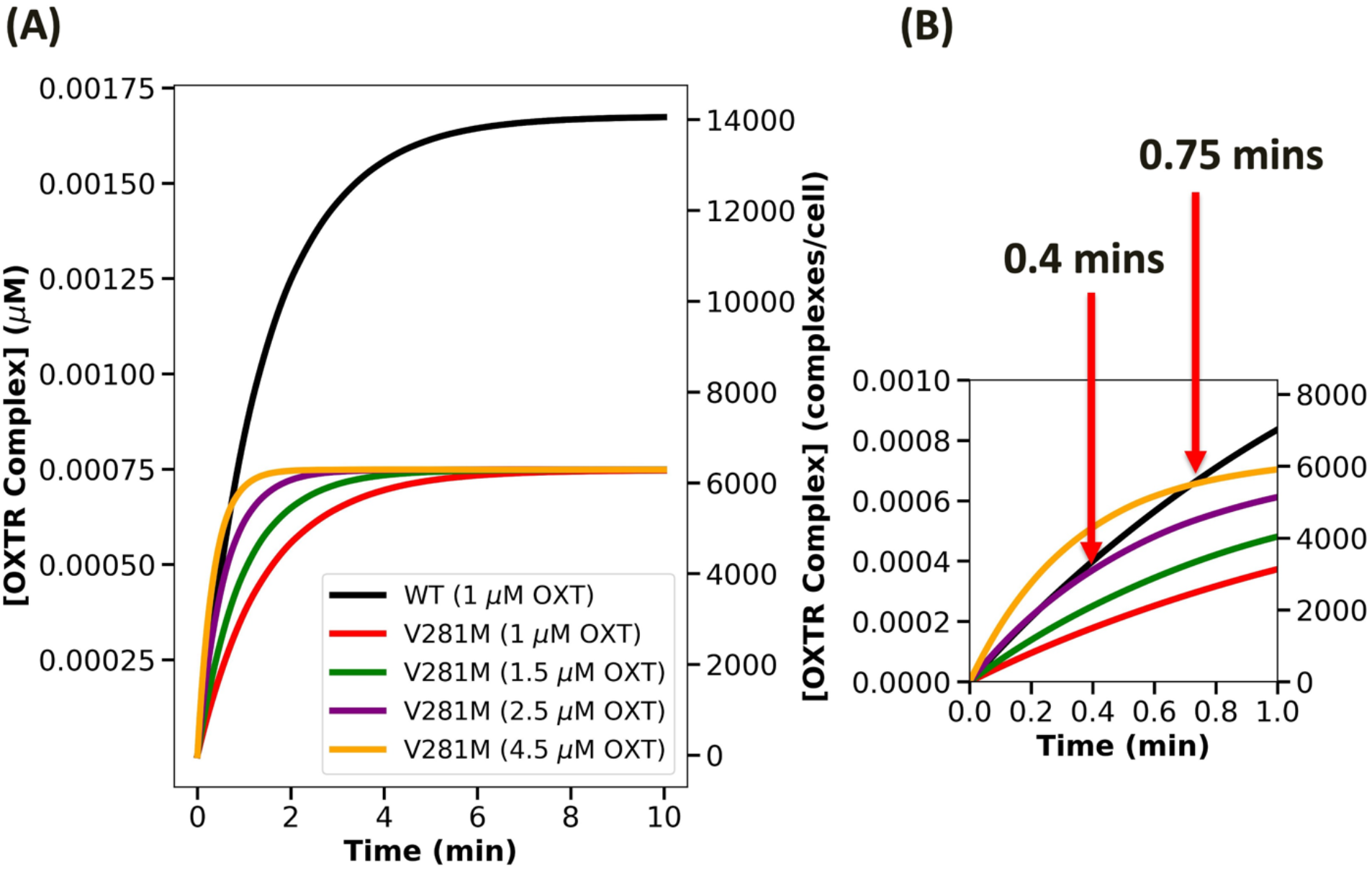
High dose of OXT can rescue the OXTRC formation for V281M OXTR in myometrial cells at early time points but not at equilibrium. **(A)** The trajectories of V281M variant complex formation with increasing doses of OXT. **(B)** The OXT dose needed to achieve the wild-type level of complex formation in 0.4 mins and in 0.75 mins.

**Figure 6:**
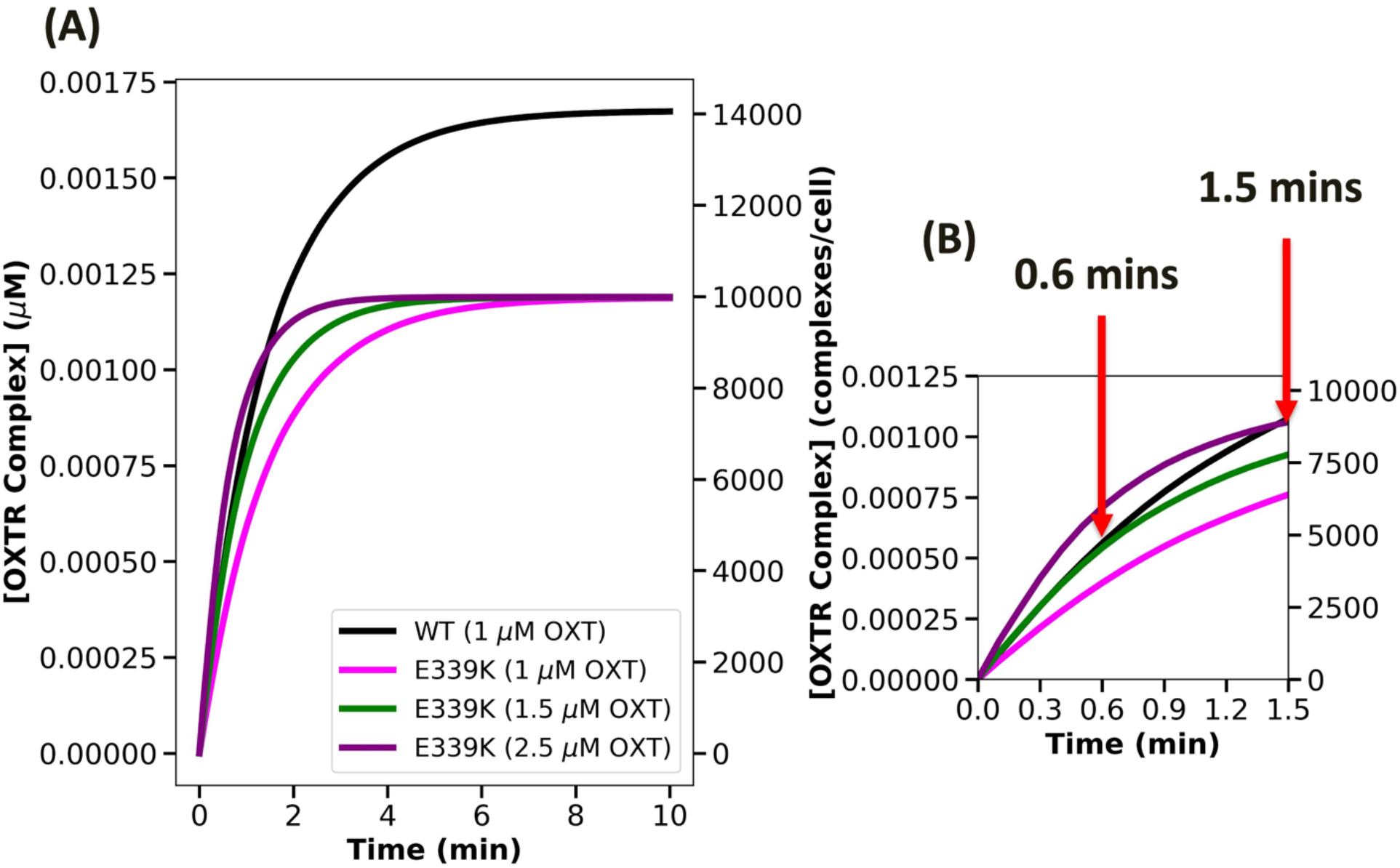
High dose of OXT can rescue the OXTRC formation for E339K OXTR in myometrial cells at early time points but not at equilibrium. **(A)** The trajectories of E339K variant complex formation with increasing doses of OXT. **(B)** The OXT dose needed to achieve the wild-type level in 0.6 mins and in 1.5 mins.

Nevertheless, the higher OXT doses can rescue OXTRC formation at early time points (within the first minute), which is a critical period for downstream signaling.^15^ Our simulation suggests that V281M OXTR can achieve the wild-type level of activation by administering 2.5 µM OXT to achieve the wild-type level of OXTRC formation until 24 seconds (0.4 min) or administering 4.5 µM OXT to exceed the wild-type level of OXTRC formation until 45 seconds (0.75 min) (**Figure 5B**). Similarly, E339K OXTRs require 1.5 µM OXT to achieve the wild-type level of OXTRC formation until 36 seconds (0.6 min), or 2.5 µM OXT to exceed the wild-type level of OXTRC formation until 90 seconds (1.5 min) (**Figure 6B**). Therefore, the OXTR-dampening genetic variants could still be adequately activated with higher OXT doses at early time points.

### Tuning the adjustment needed in binding affinities of OXTR-dampening genetic variants in myometrial cells

In addition to increasing OXT dosage, engineering OXT that binds OXTR with a strong affinity will help rescue the WT-level OXTRC formation in V281M- and E339K-carrying myometrial cells. Several OXT analogs have been engineered, with the hope of either mimicking (e.g., Pitocin) or achieving improved activation of OXTR (e.g., Carbetocin).^16^ Here, we determined the binding kinetics necessary for OXT receptor agonists to effectively bind to OXTRs and rescue WT-level OXTRC formation in V281M- and E339K-carrying myometrial cells. The binding affinity of OXT-OXTR in human myometrial cells at term is 1.6 nM (k_on_ = 6.8 × 10^5^ M^−1^ min^−1^ and k_off_ = 1.1 x 10^-3^ min^−1^). Thus, to achieve WT-level OXTRC formation within the first minute of 1 µM OXT treatment, an OXTR agonist would need to have 30 times stronger affinity (K_d_ = 0.052 nM) compared to OXT to rescue V281M OXTR and 20 times stronger affinity (K_d_ = 0.083 nM), compared to OXT to rescue E339K OXTR. Regarding the binding kinetics, the agonist would have a k_on_ = 2.28 × 10^6^ M^−1^ min^−1^ and k_off_ = 1.18 x 10^-4^ min^−1^ toward V281M OXTRs (**Figure 7A**) and k_on_ = 1.2 × 10^6^ M^−1^ min^−1^ and k_off_ = 1.0 x 10^-4^ min^−1^ toward E339K OXTRs (**Figure 7B**). Therefore, engineering small-molecule OXTR agonists to rescue early signaling of the V281M and E339K requires 20 to 30 times stronger binding than OXT.

**Figure 7:**
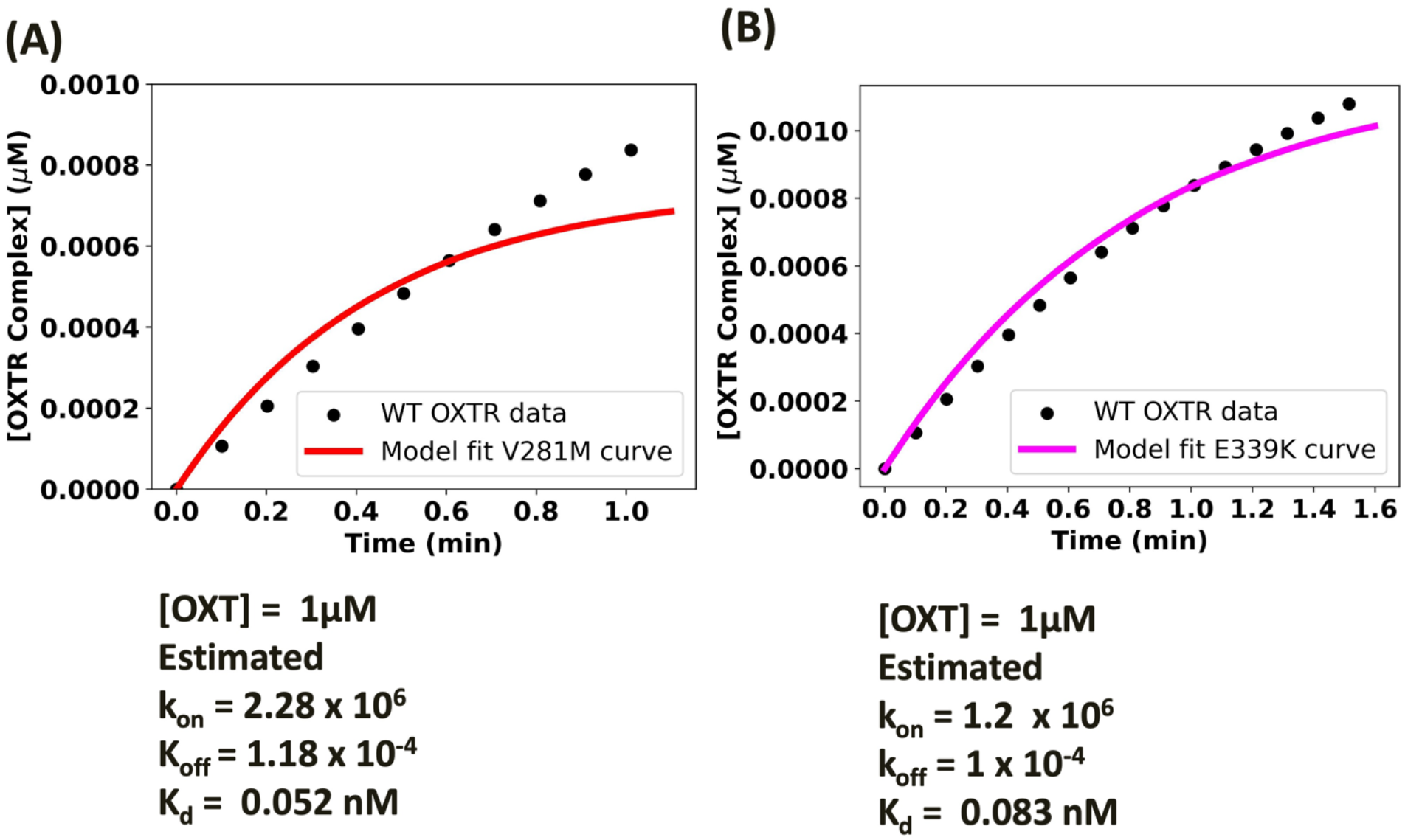
Engineering small-molecule OXTR agonists to rescue early signaling of the V281M and E339K requires 20 to 30 times stronger binding than oxytocin. **(A)** Binding affinity tuning for V281M with respect to wild type OXTRC in myometrial cells. We have estimated the association rate constant (k_on_) and dissociation rate constant (k_off_) required to achieve the level of WT OXTRC formation in 1 min by V281M OXTRC formation. The model for V281M variant (red line) was fitted to the WT OXTRC formation data (black dots). **(B)** Binding affinity tuning for E339K with respect to wild type OXTRC in myometrial cells. We have estimated the association rate constant (k_on_) and dissociation rate constant (k_off_) required to achieve the level of WT OXTRC formation in 1.5 min by E339K OXTRC formation. The model for E339K variant (pink line) was fitted to the WT OXTRC formation data (black dots).

## DISCUSSION

### Construction of a mathematical model to predict OXT responses in myometrial cells

Our mathematical model offers data-driven advancements in receptor concentrations and binding kinetics, as well as a computational advancement. The model is cell-specific through parameterization with quantitatively distinct OXTR concentrations associated with different genetic variants in HEK293T cells^5^ and human myometrial cells^10^. The specificity of this model is further bolstered by the cell-specific OXT-OXTR binding kinetics, obtained through our extensive literature review and meta-analysis. Furthermore, we arrived at the first experimentally validated (via EC_50_ experimental data prediction), mathematical model that predicts OXT–OXTR interactions in human myometrial smooth muscle cells, the cell type responsible for the uterine contractions mediating labor and delivery. This validated model leverages cell–specific OXTR expression data and OXT–OXTR binding kinetics. Importantly, our model offers three key findings: 1) differences in dosage and time to maximal occupancy between human myometrial smooth muscle cells and the heterologous expression system, HEK293T cells, 2) prediction of decreased complex formation in the V281M and E339K mutants, and 3) predictions of dosing or binding affinity necessary to potentially rescue V281M and E339K complex formation. Our model can help bridge the findings from HEK293T cells and myometrial cells. For instance, our model suggests that the OXT dose response obtained from HEK293T cells (predicted EC_50_=4.7 nM) needs to be scaled up threefold to reflect that in myometrial cells (predicted EC_50_=14.5 nM). Future adaptations to include OXTR concentrations in endogenous myometrial cells under a variety of conditions could facilitate personalized OXT dosing in clinical settings.

### Precision OXTR modeling: The power of quantitative data

Recent breakthroughs in qFlow have led to accurate, reproducible quantification and identification of differential receptor concentrations across normal and diseased cell phenotypes^17–26^. These advances enabled us to show, for the first time, how genetic variation affected the presentation of OXTR on the cell membrane. Previously, qFlow has been instrumental in measuring receptor tyrosine kinases (RTKs) that, like GPCRs, are integral to a wide array of physiological and pathological signaling pathways, regulating many facets of health and disease. Computational models have applied this receptor concentration data to offer predictions on RTK signaling. For instance, the model by Finley et al. incorporated parameters like VEGFR concentrations on the surface of healthy and diseased cells, VEGF-VEGFR binding kinetics, and distribution of VEGF-neutralizing antibodies. This model provided predictions for optimizing VEGF-targeting cancer therapies^27^. Our OXT–OXTR binding model aims to emulate the success of RTK modeling by applying quantitative OXTR data to predict improved OXT dosing for labor management.

Our study demonstrates that the tenfold difference in OXTR surface localization between the transfected HEK293T cells and transfected human myometrial cells, identified by qFlow, contributes to the three-fold difference in the EC50 of OXTR complex formation at 1 µM OXT. In other cell types, previous work has shown that differential protein localization on the cell-surface can result in differential cell response. For instance, OVCAR8, a high-grade ovarian cancer cell line, exhibits 44 times more surface AXL tyrosine kinase receptors than OVCAR3, a different ovarian cancer cell line^17^. The significant difference in AXL abundance contributes to greater resistance to anti-cancer drugs by OVCAR8.28 Overall, receptor surface localization and receptor binding affinity can be cell-specific and must be carefully considered when studying the receptor-targeted cell responses.

### Differential drug responses across and within cell types

Although transfected HEK293 cells are commonly used to study cell signaling and drug response *in vitro,*^5,29,30^ including to study of OXT–OXTR binding, this heterologous system differs from biologically relevant cells expressing endogenous proteins and can lead to differential receptor localization to the cell surface, differential activation, and differential signaling. Indeed, we have already observed a ten-fold difference in the numbers of OXTRs on the cell surface (or an over 100-fold difference in molar concentrations when taking the cell sizes into account) and a three-fold difference in binding kinetics when comparing HEK293T and myometrial cells. Our modeling showed that these differences lead to differential OXTRC formation between HEK293T cells and human myometrial cells.

### Differential drug responses across genetic variants

Genetic variants, particularly single nucleotide missense variants that are encoded in the resulting protein, could have adverse effects on GPCR protein expression and function, leading to aberrant drug responses.^31^ Missense variants in OXTR, such as V281M and E339K, were found to decrease OXTR surface localization and reduce OXT response in transfected HEK293T cells^5^. Our model shows OXTRC formation follows the same trend for myometrial cells, with a decrease of 55% for V281M and 29% for E339K compared to the wild type. The reduced OXTRC formation is due to the lower amount of cell-surface OXTRs available for OXT binding, leading to the 25% lower OXTR activation previously measured in HEK293T cells.^5^ Thus, our model can be useful for predicting the outcomes of drugs and other interventions that enhance OXTR surface localization for rescuing OXTR activation.

### New insights into OXT dosing strategies for genetic variants

We applied our model to test the OXT dose necessary to rescue WT signaling for OXTR variants V281M and E339K, which exhibited lower OXTR surface concentration compared to WT. Our model predicts that a 2.5-fold increase in OXT dose enables these variants to reach or surpass the wild-type binding level within the first 20 seconds of OXT administration. This finding is significant because key cellular responses, such as calcium mobilization, are initiated within this timeframe^15^. However, despite the ability to increase OXT concentration to rescue the V281M and E339K variants in the short term, they cannot attain the maximal binding level of wild-type OXTR. This paradox between short-term (seconds) activation rescue and long-term (several hours) equilibrium deficiency suggests two needs for OXT dosing strategies. First, a need for additional modeling of the full OXTR signaling cascade to determine how the V281M and E339K signal propagates and on what timeframe. Second, experimental studies are needed of the 2.5-fold higher OXT rescue dose, determined by the model, to see if signaling is indeed rescued in the V281M and E339K mutants.

### The OXT-OXTR binding affinity can be altered in different biological contexts

Our model suggests that a 20-to 30-times increase in binding affinity can rescue the time to WT-level OXTRC formation in V281M and E339K cells—raising the question of whether it is possible to achieve such a significant increase in binding affinity. Human myometrial OXTRs bind more strongly to OXT at term compared to the ‘non-pregnant’ and ‘1st trimester’ myometrial OXTRs (**Table 2**).^32^ While our model employs the binding parameters from myometrial cells at term to predict the OXT dose response during labor, future modeling can probe downstream signaling differences at term versus non-pregnant conditions. In addition, cholesterol and magnesium modulate the binding affinity of OXTR. Cholesterol acts as an allosteric modulator on OXTR to induce a high-affinity state for ligands,^33^ and depleting cholesterol resulted in a 110 times lower binding affinity.^34^ Magnesium forms a coordination complex with OXT and OXTR, and the omission of magnesium can decrease the binding affinity by about 1500-fold.^35^

Although the depletion of magnesium or cholesterol decreases OXT binding affinity and cannot be applied to rescue deficient V281M and E339K activation, it shows the possibility of manipulating OXT binding affinity. Alternative biologically relevant approaches include the engineering of small-molecule OXTR agonists with higher affinity for variant OXTRs. Current engineered agonists, such as carbetocin ^36^ and OXT analogs, including peptide, non-peptide, and hybrid peptide-small molecules,^16,37,38^ all have equivalent or only slightly weaker OXTR binding affinities compared to OXT. With the need for rescuing OXTR activation in OXTR-deficient cells, the engineering of small molecules with enhanced OXTR binding affinity could be pursued.

### Advancing a biochemical signaling framework for the OXTR modeling field

Several types of GPCR models have previously been developed particularly for pharmacology^39,40^, many recent models have focused on identifying ligands that can induce differential signaling through the same receptor (biased agonism)^41,42^. Furthermore, GPCR modeling has recently focused on delineating signaling kinetics, since this can be used to predict downstream activation^43,44^. As such, our meta-analysis of binding kinetics is a first step towards developing a more detailed signaling model for OXT-OXTR.

Although other OXTR signaling models, including those that involve kinetic studies, have been developed, their limitations prevent using those models in dosage strategies. For example, a mechanistic systems biology model predicted that the A218T OXTR genetic variant affects downstream events by altering receptor stability, activation, and signaling^45^. However, a limitation of that model is the assumption of OXTR concentrations and binding kinetics based on the data from a non-OXTR GPCR (i.e., serotonin receptor) in HEK293 cells. These parameters are substantially larger than our OXTR-specific parameters. The assumed OXTR concentration is 5.5 times higher than our qFlow OXTR data, the assumed k_on_ is ∼6.8 -fold higher, the assumed k_off_ is 30,000 times higher, and the resulting Kd is 4464 times higher than the parameters used in our model. Although these model types are powerful, they fall short of providing a detailed understanding regarding the variations in OXTR cell surface presentation, binding, and function that differ among cell types and genetic variants. Our work fills this gap by integrating and simulating the biochemistry of OXT signaling, and then applying this model to quantify how genetic variants alter signaling. Additional needs to move this model towards predicting the OXTR signaling cascade would require additional data on the OXTR trafficking dynamics (e.g., OXTR internalization, recycling, and degradation rates) and signal transduction processes (e.g., G protein activation, GRK-mediated phosphorylation, and β-arrestin binding.), which in many cases are limited.

### Conclusion

This work presents the first mathematical model predicting OXT-OXTR binding dynamics. This model, parameterized through a comprehensive literature search and advanced quantitative flow cytometry (qFlow) measurements, narrows the knowledge gap between experimental findings in widely utilized HEK293T cells and the lesser-studied biologically relevant OXTR-expressing cells, myometrial smooth muscle cells. Our model achieves the necessary validation by successfully replicating the experimental results of OXT-OXTR interactions in HEK293T cells and, crucially, it predicts the optimal OXT doses to potentially rescue deficient signaling in OXTR genetic variants.

This model delivers vital insights into OXT-OXTR binding dynamics in myometrial cells where experimental data is scarce. Furthermore, the model demonstrates immense potential for refining OXT dosing strategies in labor induction and augmentation. The recommended OXT dosage for labor induction and augmentation varies considerably, with a difference of up to 20-fold ^4–6,14^. Higher OXT doses carry risks like uterine rupture and hyperstimulation, while lower doses may prolong labor and increase the likelihood of fetal infection and cesarean delivery ^5–9^. Our model has the potential to determine the minimal effective OXT dose for pregnant women with impaired OXT-OXTR binding capacity, whether due to environmental or genetic factors, thereby achieving the desired wild-type binding level. Given the complexities of OXT dosing in these clinical scenarios, our model offers a path towards more accurate and safer therapeutic approaches, customizing treatments to meet individual requirements while reducing the risks associated with imprecise dosing.

## Methods

### Literature mining for OXT receptor binding affinity data

#### Databases used to perform the literature search

We performed an extensive literature survey using several databases, including *Google Scholar*, *PubMed, LitSense, BioModels, B10NUMB3R5,* and *UW Library.* We used the following keywords to perform the search: “OXT”, “OXT receptor”, “binding affinity”, “myometrial cells”, “HEK cells”, “equilibrium dissociation constant”, “K_d_”, “association rate constant”, and “dissociation rate constant”.

#### Eligibility criteria

We included *ex vivo* studies that measured the binding affinities of OXTR in myometrial cells from human (both term and non-term women) or nonhuman animals including rat, sheep, cattle, and guinea pig. We also included *in vitro* studies that measured the binding affinities of OXTR-transfected HEK293T cells. We included studies that measured the binding affinity of OXT to OXTR only. We excluded the studies that measured the binding affinity of other GPCRs than OXTR. We excluded studies that measured OXT–OXTR binding affinity in human breast carcinoma cells. Additionally, we omitted the studies that used cloned human OXTR expressed on COS-1 cells or studied OXTR on rat adipocytes. We have also excluded those studies that measured the binding affinity of antagonists to OXTR.

#### Data extraction

We used Image J^46^ to extract data from Phaneuf et al.^11^ to quantify it for our analysis. We used the following formula to calculate the reaction rate from time-course of OXTR binding data: k = Ln(2)/t_1/2_, where k is the rate constant of reaction, t_1/2_ is the time to reach 50% maximal binding, and Ln(2) is the natural logarithm of 2. We used the association and dissociation rate constants for HEK293T cells provided by Gulliver et al.^12^

### Data analysis using the Bayesian hierarchical model

We employed a Bayesian hierarchical model with linear regression and varying intercepts to analyze the equilibrium dissociation constant (K_d_), which is a measure of binding affinity. Bayesian hierarchal models with linear regression and varying intercepts allow us to effectively analyze complex datasets, account for hierarchical structure and heterogeneity, and improve parameter estimation. The model was implemented using the PyMC3^47^ library in Python.

### Bayesian hierarchical model structure

In this hierarchical model with linear regression and varying intercepts, each group (in this case, each cell type assayed) has its own baseline binding affinity (intercept), reflecting differences in the binding affinity between cell types. However, the effect of the predictor variable (the method employed) on the dissociation rate constant (K_d_) is assumed to be consistent across all groups, indicating a fixed slope. This means that while the baseline binding affinity (intercept) can vary between cell types assayed, the influence of the method employed (slope) on the binding affinity (K_d_) remains constant. Therefore, regardless of the cell type assayed, a change in method employed will produce proportional change in the binding affinity.

### Bayesian hierarchical model parameters

The Bayesian model parameters were defined as follows:

### Hyperpriors

We used a normal prior distribution for a parameter “μ_0_” with a mean equal to the logarithm of the mean binding affinity and a standard deviation of 1. We also defined a half-Cauchy prior distribution for a parameter “σ_0_” with a rate parameter of 1.

### Varying intercepts

We defined a normal prior distribution for a group-level parameter named “source_intercept.” This prior distribution has a mean equal to the value of the “ μ_0_ “ parameter and a standard deviation equal to the value of the “σ_0_” parameter.

### Common slope

We defined a normal prior distribution for a parameter “β” with mean 0 and standard deviation 1.

### Bayesian model error

We defined a half-Cauchy prior distribution for a parameter named “σ” with a rate parameter of 1.

### Data likelihood

The likelihood of the observed data was defined using a normal distribution. The mean of this distribution was set to the expected value theta, and its standard deviation was set to the value of the “ σ “ parameter. The observed data were specified using log-transformed K_d_ data. The data were then reverted to their original form for plozng and analysis.

### Data observation

The observed data collected, equilibrium dissociation constants (K_d_) from various cell lines, including myometrial smooth muscle cells from different species and HEK cells. We performed the meta-analysis on all the cell lines and then repeated the analysis, restricting our observed data to human cell lines.

### OXT receptor measurements

#### Transfected HEK293T cells

Plasma membrane surface OXTR concentrations on transfected HEK293T cells were taken from published quantitative flow cytometry data^5^. We converted the number of membrane receptors/cell to molarity, using the following equation:

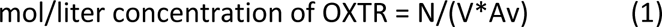

where, N = the number of membrane receptors/cell, V = volume of the HEK293 cell, and Av = Avogadro’s number. The volume of a HEK293 cell is given as 1046 cubic micrometers (1.046 × 10^−12^ liter)^48^.

#### Transfected hTERT human myometrial smooth muscle cells

Plasma membrane surface OXTR concentrations on transfected myometrial cells were taken from our recently published data that were collected using qFlow^10^. We used a procedure similar to that described above to convert the concentrations to molarity. We assumed a myometrial cell volume of 14,047 cubic micrometers (1.4 × 10^−11^ liter) ^49^.

### Mathematical and Computational Methods

#### Model construction

We constructed a mathematical model to elucidate the OXT binding of WT OXTR and five OXTR variants (V281M, E339K, V45L, P108A, and L206V) as shown in the schematic of the model. A schematic of variants and their positions in the largest OXTR crystal structure is shown in **Figure 1**. We assumed that the ligand OXT binds to the OXTR on the plasma membrane and described the binding using ordinary differential equations as follows:

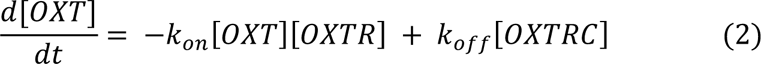

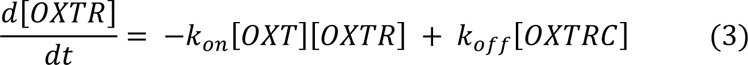

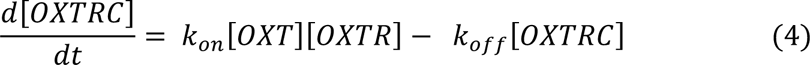

where [OXT] is the concentration of the OXT ligand, [OXTR] is the concentration of OXT receptor, and [OXTRC] is the concentration of OXT receptor complex. The association rate constant of the chemical reactions is given by k_on_ and dissociation rate constant is given by k_off_.

**Equation 1** describes the association and dissociation of OXT ligand over time, **Equation 2** describes the association and dissociation of OXT receptor over time, and **Equation 3** describes the association and dissociation of OXT receptor complex over time.

#### Model assumptions

We assumed that the process of OXT and OXT receptor binding on the plasma membrane is continuous. We assumed the binding kinetics is not affected by the studied variants because they are in the OXTR non-binding sites. We assumed that there is no clearance of OXT and that there is no endogenous OXT in this model. We also assumed that there is no desensitization of the receptor binding.

#### Model parameterization

We performed an extensive literature survey to parameterize the model. We extracted the data from the literature and calculated an association rate constant (k_on_) and a dissociation rate constant (k_off_) for human myometrial cells from the given equilibrium dissociation constant (K_d_, binding affinity) in Phaneuf et al. ^11^. The results were as follows: k_on_ = 6.8 × 10^−4^ nM^−1^ min^−1^ and k_off_ = 0.0011 min^−1^ (K_d_ = 1.6 nM) for transfected human myometrial cells. For transfected HEK293 cells, we used published association and dissociation rate constants from Gulliver et al. ^12^ and the results were as follows: k_on_ = 8.8 × 10^−3^ nM^−1^ min^−1^ and k_off_ = 0.005 min^−1^ (K_d_ = 0.56 nM) for OXTR-transfected HEK293 cells. We chose this study because the kinetics were measured by radioligand binding, which aligns with our choice for myometrial K_d_ value from Phaneuf et al., which was also measured by radioligand binding, thus minimizing the method-associated variation in K_d_ measurements. The binding affinities from different sources and experimental processes for different cell types are shown in **Figure 2**.

#### Model simulations

We performed simulations of the model using the model parameters given above and initial conditions given in **Table 2**. We used the Scipy, Numpy, and Matplotlib libraries of Python 3.9.7 to perform simulations, and for parameter estimation we used the lmfit package (<u>Levenberg-</u> <u>Marquardt</u> method from <u>scipy.optimize.leastsq)</u>. The detailed Python script will be provided on request.

## Author Contributions

**P.D.** led computational modeling methodology, construction, data curation, and simulation; conceptual validation, analyses, visualization, and writing – reviewing and editing. **X.L.,** and **A.S.** data curation. **K.L.T.** data curation, meta-analysis, and visualization (Figure 2) **Y.F.** led writing – original draft, led writing – review and editing, biological and experimental data investigation and interpretation; computational modeling validation, analyses, visualization, project administration for clinical and translational collaboration, and supervision of data curation, protein structure and kinetics visualization, meta-analysis. **S.K.** protein structure and binding kinetics methodology and visualization (Figure 1). **A.F., E.R.,** and **M.M.** clinical and translational interpretation and validation of data curation and model analyses, writing – review and editing. **S.K.E.** clinical and translational interpretation and validation of data curation and model analyses, conceptualization-model for OXTR, data curation, funding acquisition, project administration, resources, supervision, writing – review and editing. **P.I.I.** conceptualization-model for OXTR, theory, and scope; data curation; validation methodology; analyses; investigation; methodology; project administration; resources; funding acquisition; supervision of modeling, data-curation, meta-analysis, visualization, and writing; writing – original draft, and writing – review and editing.

## Acknowledgements

This research is supported by the National Institute of Child Health and Human Development (R01 HD096737). The authors declare no competing financial interest.

